# Coarse-Grained Model of the SNARE Complex Shows that Quick Zippering Requires Partial Assembly

**DOI:** 10.1101/294181

**Authors:** N. Fortoul, M. Bykhovskaia, A. Jagota

## Abstract

Neuronal transmitters are released from nerve terminals via the fusion of synaptic vesicles with the presynaptic membrane. Vesicles become attached to the membrane via the SNARE complex. The SNARE complex comprises the vesicle associate protein Synaptobrevin (Syb), the membrane associated protein syntaxin (Syx), and the cytosolic protein SNAP25, which together form a four-helical bundle. The full assembly of Syb onto the core SNARE bundle promotes vesicle fusion. We investigated SNARE assembly using a coarse-grained model of the SNARE complex. The model retains chemical specificity, and was calibrated using single molecule experiments and all-atom molecular dynamics simulations. Steered force-control simulations of SNARE unzippering by peeling off Syb were used to set up initial disassembled states of the SNARE complex. From these states, the assembly process was simulated. We found that if Syb is in helical form, then the SNARE complex assembles rapidly, on a sub-microsecond time-scale. We found that assembly times grow exponentially with separation distance between Syb and Syx C-termni. The formation of helical turns is likely to substantially decelerate the assembly, consistent with single molecule force experiments that show SNARE assembly duration on the time-scale of hundreds of ms. Since synaptic vesicle fusion occurs at a sub-millisecond time-scale, our results indicate that for biologically relevant rapid assembly the SNARE complex needs to be partially zippered and its constituent helices brought into proximity, possibly by means of molecular chaperones.

## 1. INTRODUCTION

Neuronal transmission occurs via fusion of synaptic vesicles which are docked at the synaptic membrane (1). The SNARE (soluble N-ethylmaleimide-sensitive factor attachment protein receptor) (2, 3) proteins are predominantly responsible for providing the adhesive force to overcome the electrostatic and hydration repulsion between the vesicle and the membrane. SNARE proteins form a four-helical bundle comprising the vesicle bound Synaptobrevin (Syb or v-SNARE) and the plasma membrane bound t-SNARE which includes Syntaxin (Syx) and SNAP-25. The latter protein comprises two helical domains (SN1) and (SN2) (4, 5). Zippering of t-SNARE and v-SNARE is thought to provide adhesive forces and energies to overcome the repulsion between the vesicle and the membrane.

The atomic structure of the SNARE bundle has been determined by x-ray crystallography (6); it comprises sixteen layers, and their contributions to the overall SNARE assembly have broadly been identified (7, 8). However, the mechanistic details of SNARE assembly kinetics and its effect on the fusion process are still obscure. The disassembly/assembly pathway of the SNARE complex has been investigated employing optical and magnetic tweezers (9, 10), as well as by molecular dynamics simulations, including all-atom (AA) (11), and coarse-grained (CG) approaches (12). Using optical tweezers to pull at the C-termini of Syb and Syx, Gao et al. (9) studied SNARE disassembly/assembly *in vitro*, producing accompanying force-displacement relationships. In particular, Gao et al. (9) determined that SNARE disassembly takes place in stages that are accompanied by melting of Syb, and found the distribution of probabilities for Syb helix melting/re-formation as a function of applied force. These SNARE unzippering/zippering stages were confirmed by a magnetic tweezers study (10). This study found that there are significant activation barriers to SNARE assembly when studied *in vitro* with characteristic assembly rate of hundreds of ms (zero-force C-terminal-half zippering assembly rate was found to be 5.93 s^-1^). These relatively slow assembly times are not consistent with the rapid process of action potential evoked fusion *in vivo*, which occurs at a sub-millisecond time-scale (2). This suggests the hypothesis that the large *in vitro* versus *in vivo* discrepancy in assembly time is due to some agent pre-forming helical structures of Syb and Syx, producing a partially assembled bundle. A testable consequence of this hypothesis is that Syb should assemble rapidly if it is in a helical form and partially assembled onto t-SNARE.

We test this hypothesis by performing CG molecular simulations of SNARE assembly. We have previously developed a CG model of the SNARE complex where each residue was represented by a single bead (12). The model was calibrated by experimental and AA simulation results and combined with a continuum model of the synaptic vesicle and membrane to determine the number of SNAREs required for synaptic vesicle to membrane docking. More recently, a four residue/bead CG was developed (13) to study the cooperativity and self-organization of the SNARE complex. This model took advantage of the energy landscape obtained in optical tweezers experiments (9). Here we further developed our CG model to simulate unzippering and reassembly of the SNARE bundle. The model uses a single bead per residue, thus retaining chemical specificity while easily accessing microsecond timescales. In order to better interpret CG simulation results, we also present an analytical scaling model for the assembly process. Our results indicate that if helical Syb is brought into proximity of t-SNARE, assembly occurs within biologically relevant times, on a sub-microsecond scale, suggesting that the discrepancy between *in vitro* and *in vivo* time scales is due to the presence of molecular chaperones that serve as a template in the latter case.

## 2. MATERIALS AND METHODS

The model: CG-MD simulations of the SNARE bundle were conducted using Brownian dynamics (BD) (12, 14). The CG model of SNARE, shown in Fig. 1, is protein-residue based, each one being represented by a bead at the location of its alpha-carbon.

**FIGURE 1.**
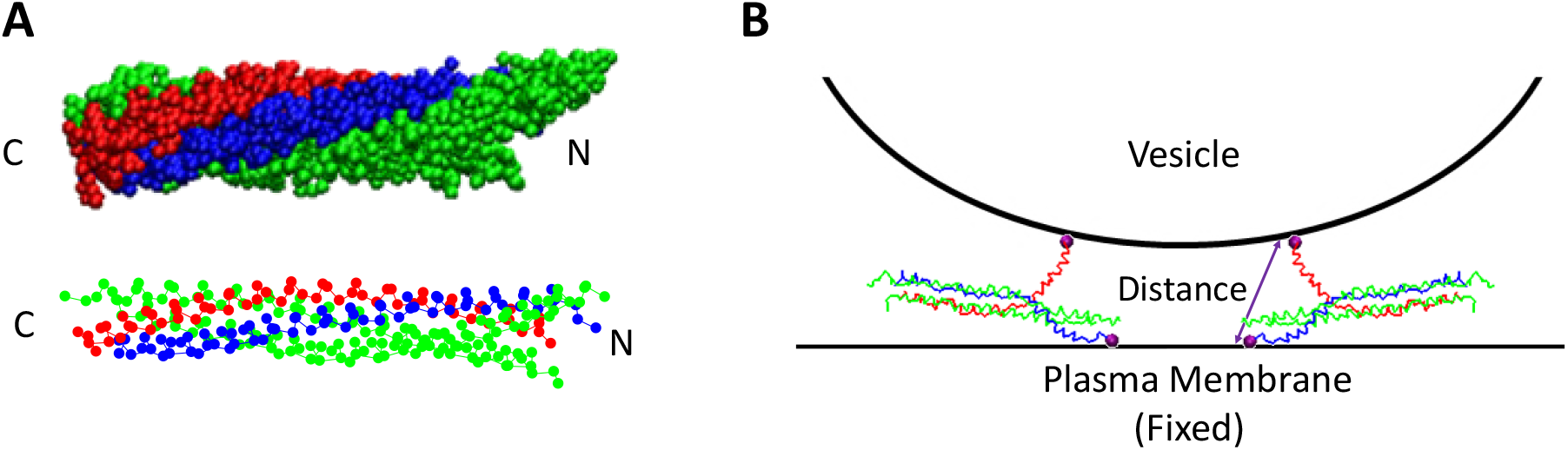
(*A*) The all atom structure of SNARE is shown with each atom being represented by a single bead. The corresponding CG structure of SNARE is also shown with each bead representing a residue and located at the position of its alpha carbon. The SNARE structures include Syb (*red*), Syx (*blue*), SN1 (*green*), and SN2 (*green*) and the C and N termini are indicated. (*B*) A schematic in which the synaptic vesicle and plasma membrane are shown along with the CG model of SNARE. The pulling beads are at the C-termini of Syb and Syx (*purple*). The Syx pulling bead is held fixed while a force is applied onto the Syb pulling bead.

The CG model of SNARE and the forcefield used was based on the model described in Fortoul et al. (12) where an ENM (Elastic Network Model) was used to hold the shape of helices and MJ (Miyazawa-Jernighan) potentials were used for helix-helix interactions. The majority of the simulations conducted in our previous work (12) were done at quasi-equilibrium conditions at 0 K, and the simulations conducted in this paper were done at 300 K, so some adjustment in parameters was required.

An ENM (15, 16) was used to represent intrahelical interactions and hold the shape of the helices. In this method, all beads within a single helix are connected to all other beads in the same helix within a cutoff distance, *R_c_*, using harmonic springs and the energy potential

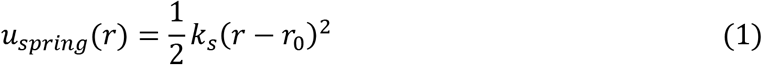

where *k*_*s*_ is the spring constant, *r* is the distance between 2 beads, and *r*_0_ is the natural length of the spring. In Fortoul et al. (12), the value of *k*_*s*_ used was 0.0963 N/m in order to match single helix all atom fluctuations to single helix coarse-grained fluctuations. However, at 300 K slightly stronger springs were required to hold the helical shape. In order to determine an appropriate value of *k*_*s*_ several simulations of the CG structure of the SNARE bundle were conducted for 20 ns using different values of *k*_*s*_. These were compared to all-atom simulations of SNARE; in particular we compared fluctuations of the alpha-carbons and found an RMSD of 4.7 A to 17 A for ks in the range 0.0963 N/m 0.1926 N/m (12). We chose a value of 0.1685 N/m for *k*_*s*_ because this was the minimum value that maintained a stable helical structures and was within this acceptable range.

To increase simulation speed, we also adjusted the value of cut-off distance for ENM springs. This has a significant effect of computational speed because a smaller value of *R*_*c*_ creates a smaller network of springs and requires fewer calculations. In Fortoul et al. (12) a value of 20 A was used. To find a minimum value of *R*_*c*_ that could be used without compromising simulation validity, several 20 ns CG simulations of SNARE were completed using different values of *R*_*c*_. *R*_*c*_ could be decreased to a minimum of 16 A without compromising the SNARE structure (using a *k*_*s*_ of 0.1685 N/m); therefore the value was decreased to 16 A.

Interhelical interactions were modeled using Miyazawa and Jernigan (MJ) energies (17–19) that provide chemical specificity. For a particular bead, every bead within the cutoff distance, *R*_*c_MJ*_, of that bead (not within the same helix) has an interaction with that bead. For each interaction type there is a contact energy, *e*_*ij*_, that is dependent on the two interacting bead types, *i* and *j* (e.g. Leu-Leu vs. Leu-Trp). The value of *e*_*ij*_ is adjusted for a particular system as

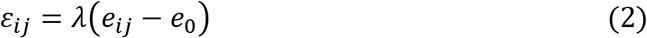

where *ε*_*ij*_ is the scaled contact energy, *λ* is a scaling parameter, and *e*_0_ is a shifting parameter. *λ* and *e*_0_ are the same for all interaction types. A negative value of *ε*_*ij*_ represents an attractive interaction, while a positive value of *ε*_*ij*_ represents a repulsive interaction. Following (20), interaction energies are implemented as interaction potentials that depend on both the sign of *ε*_*ij*_ as well as if the distance between bead is larger or small than the distance at the potential minimum, 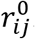.

The attractive interaction potential (*ε*_*ij*_ < 0) is

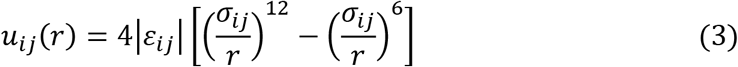

The repulsive interaction potential (*ε*_*ij*_ > 0) when 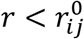 is

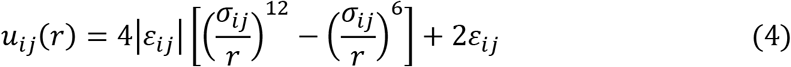

The repulsive interaction potential (*ε*_*ij*_ > 0) when 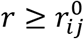 is

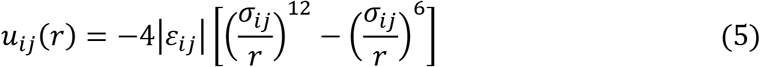

where *r* is the distance between the two beads and *σ*_*ij*_ is the interaction radius. *σ*_*ij*_ is the scaled average of the Van der Waals (VDW) diameters of the two bead types:

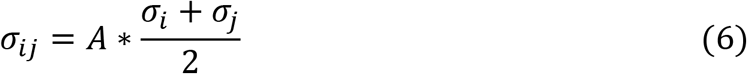

where *A* is the scaling parameter, *σ*_*i*_ is the VDW diameter of bead type *i*, and *σ*_*j*_ is the VDW diameter of bead type *j*. In order to ensure that the proper parameters were being used, several 20 ns relaxation simulations of the SNARE bundle were conducted using different values of *λ*, *e*_0_, and *A*. Parameters were chosen to produce the right SNARE bundle width and overall bundle shape compared to that of the AA relaxed crystal structure. *λ* was adjusted from 0.30 in our previous study to 0.37; *e*_0_ was adjusted from 0 in our previous study to 0.95*kT; *A* was kept at 0.8 from the previous study.

In addition to CG simulations in which Syb remained helical, two additional cases were studied (see Supporting Material). In these simulations Syb was allowed to “melt” following a force-dependent Monte Carlo (MC) scheme based on the optical tweezers experiments reported by Gao et al. (9). A critical enhancement of the CG model is the ability to capture transitions between helical and random coil states by removal or reinstatement of the elastic network bonds (the backbone is always retained), based on single-molecule stretching experimental data. During subsequent assembly, Syb was returned to helical form either rapidly upon release of force or using a criterion based on distance from t-SNARE. The purpose of these additional simulations was to judge the robustness of our conclusions to variations in how Syb forms the helix. Further details on these models are provided in Supporting Material.

In all simulations performed the initial configuration was determined using the x-ray structure 1N7S (7, 11, 12) which included Synaptobrevin (residues 27-88) (Syb), Syntaxin-1A (residues 191-256), SNAP25 (residues 7-83 of the SN1 fragment and residues 141-204 of the SN2 fragment). This structure was simulated for 40 ns using AA MD in the NAMD software package (11). The corresponding CG structure was generated as described above and relaxed for 20 ns. Subsequently, a fixed bead was attached to the C-terminal bead of Syx, and a second fixed bead was attached to the C-terminal bead of Syb. The SNARE structure with the two fixed beads was then relaxed using BD for another 20 ns. All subsequent CG simulations were performed using the initial configuration determined by the final timestep of the 20 ns BD run with fixed beads.

### 2.1. First Step: Separating Syb from the SNARE Bundle Under Force Control

Steered dynamics under force control (FC) simulations were used to unzipper the SNARE bundle to set up initial states for SNARE assembly studies. A fixed bead was attached to the C-terminal end of Syx and a pulling bead was attached to the C-terminal end of Syb, as shown in Fig. 1, with an applied force of 0.056 pN/ns along the vector connecting the two beads. This forcing rate was chosen by running simulations with decreasing forcing rate until it was slow enough that the system was close to equilibrium as determined by agreement between mean forces on the two pulling beads (Supporting Material, Fig. S3). In addition to the case where Syb remained helical, we also simulated the first step while allowing Syb to melt following a force-dependent criterion described in Supporting Material.

### 2.2. Second Step: Re-assembly of SNARE

After an initial state was established using FC, assembly simulations were performed by releasing the force on the Syb pulling bead and allowing the SNARE to relax and re-assemble. Reassembly of the SNARE bundle was studied from a variety of SNARE initial states ranging from very slightly unzippered to fully unzippered SNARE. For the cases where Syb was allowed to melt in the first step, two re-assembly schemes were simulated. In the first one, when the force is released on the Syb pulling bead, all ENM network springs are replaced which quickly reforms the Syb helix. This is meant to test if it matters whether the helical state is preceded by a melted random coil state for Syb. In the second scheme, we employed a distance criterion such that, as alpha carbons get within a certain distance of the SNARE bundle, their helicity is reinstated. This was to test if delayed helix formation of Syb affected our conclusions.

## 3. Results and Discussion

### 3.1. Syb in helical form assembles onto the core t-SNARE within hundreds of nanoseconds

The initial state prior to assembly (step 1) was produced by unzippering of Syb from SNARE under FC, with Syb in a fully helical state. FC on the Syb C-terminal bead was used to completely pull the SNARE bundle apart to where all layers are unzippered (a distance of ~20 nm between the Syb and Syx C-termini) as shown in Figs. 2 *A* and *B* (blue traces). Relaxation to assembly (step 2) followed release of the force with various values of initial opening.

**FIGURE 2.**
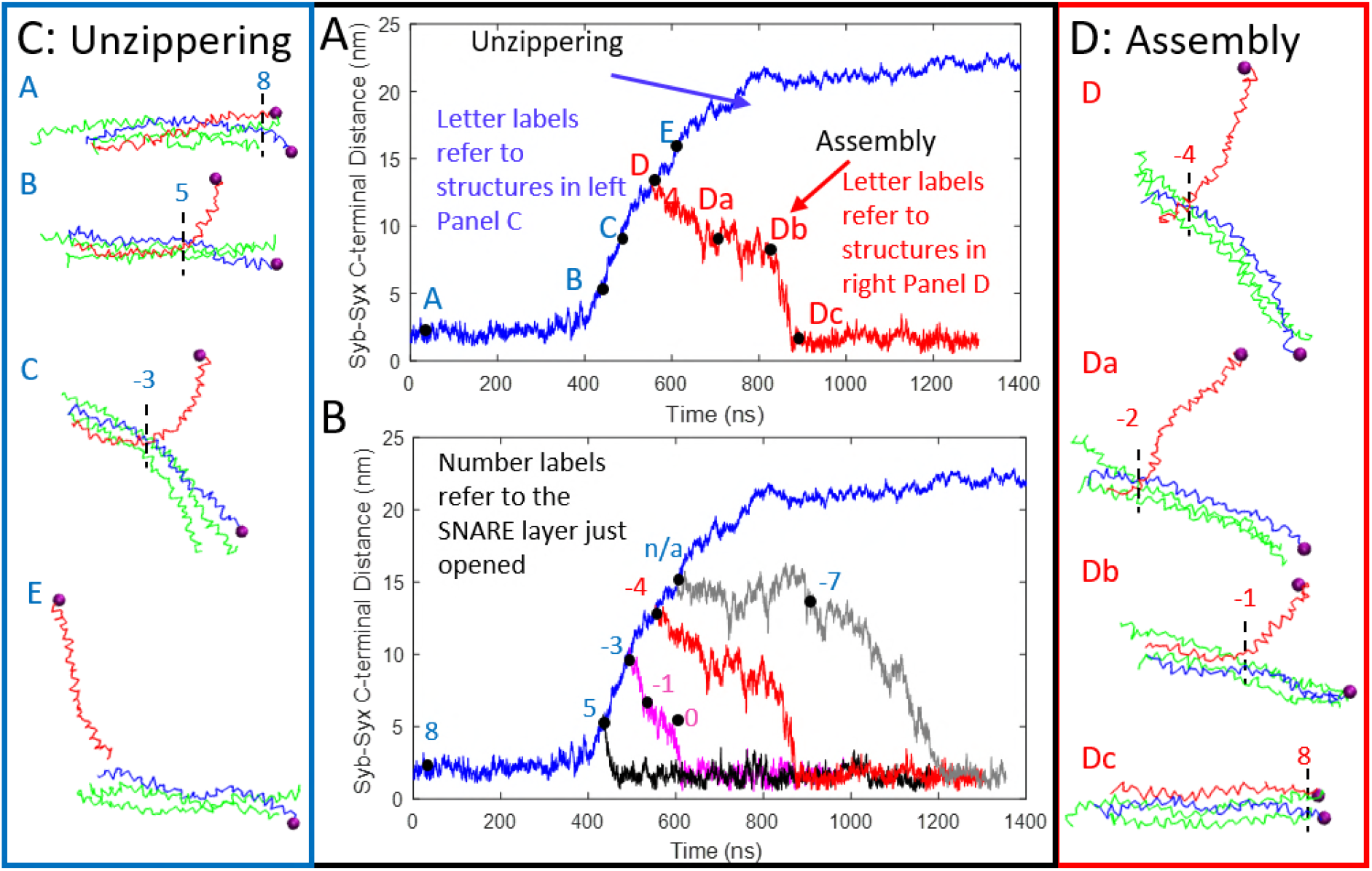
(*A*) Syb-Syx C-terminal distance as a function of time (ns) for unzippering of Syb in fully helical form off SNARE (*blue trace*). Also shown is an example of an assembly simulation (*red trace*). The unzippering trace is labeled by letters corresponding to snapshots from the simulation trajectory shown in (*C*). The assembly trace is labeled by letters corresponding to snapshots shown in (*D*). (*B*) The force was released and the assembly process begun at several points within the unzippering trajectory (e.g., *black, magenta, red, and gray traces)*. Labels refer to the layer up to which the SNARE bundle is zippered; see also (*C*) and (*D*). For small initial SNARE openings of only a couple of layers (*black*) the bundle assembles at a relatively constant fast rate. For intermediate initial openings (*magenta and red*) the bundle starts to assemble initially at a slow rate and then assembles at a faster rate. (*C*) Snapshots of SNARE during unzippering. (*D*) Snapshots (*D-Dc*) for one specific assembly case. The bundle assembles slowly from layer −4 to layer −1 (near the ionic layer). Then from layer −1 to layer 8 it assembles quickly. For large initial openings, where no layers are initially in contact (*gray trace in (B)*), Syb fluctuates at roughly a constant distance from the bundle until layer −7 is assembled, then the rest of the bundle assembles at an increasing rate. (Only a few of the BD assembly simulations are shown in this figure for clarity.)

At several points during the unzippering trajectory, the force on the Syb pulling bead was released, and the SNARE bundle allowed to relax. For example, point D Fig. 2 *A* indicates the initial state for a relaxation run beginning at Syb-Syx unzippered to layer −3 (13.4 nm). A fully assembled SNARE bundle is indicated by a Syb-Syx distance of ~2 nm and is zippered to layer 8 which is about its value in a relaxed and fully assembled SNARE bundle taken from the x-ray crystal structure. Fig. 2 *B* shows how, as the starting Syb-Syx separation increases, so does the time required for assembly. For small initial separations of up to layer −3 (~10nm), the SNARE bundle immediately begins to assemble at a relatively constant rate when the applied force is released, and completes assembly in 250 ns or less. For SNARE unzippered to layers −3 and −4 (initial separations of ~10nm to ~13.5 nm) the bundle assembles in two major steps. The bundle initially begins to assemble at a relatively slow rate until the bundle is zippered up to layer −1 after which the rate of assembly increases; see the run initially unzippered to layer −4 (13.4 nm) in Fig. 2 *B* and *D*. Layer −1 is very close to the ionic layer (layer 0), a particularly adhesive region within SNARE, which is known to help hold SNARE in a partially zippered state (9). Snapshots are shown in Fig. 2 *D* for one of the assembly trajectories. For SNARE unzippered past layer −4 (separations >13.4nm), there are again two distinct regions during relaxation. The first region is more of a plateau in which the distance of Syb from the bundle, on average, changes little. At some point in time (see grey trace in Fig. 2 *B*), there is a transition as Syb initiates contact with the SNARE bundle (at about layer −7 in this instance). Subsequently, Syb increases its contact with the rest of the SNARE bundle. For even greater initial separation, there is no reassembly within the duration of the simulation. For the cases where assembly is initiated, it runs to completion, and the final re-assembled structure is essentially the same as the starting one, as described later in more detail. That is, a helical Syb assembles in about the microsecond time scale if is in partial contact with t-SNARE.

### 3.2 Comparison of Initial and Final SNARE Bundle Structures

An important question is: does the re-assembled SNARE match the initial SNARE structure? The reassembled structures are overlaid on the initial SNARE bundle structures in Fig. 3. Characteristics of the ensemble of reassembled states, such as its width and size, are very similar to those of the initial state. Some of the SNARE helices have shifted slightly around the bundle. Fig. 3 *A* shows root mean square difference (RMSD) 5.5 ± 0.6 A, representing thermal fluctuations in the starting ensemble of SNARE structures over a 1 µs simulation. Fig. 3 *B* shows the average RMSD of 11.1 ± 1.6 A between the ensemble of reassembled SNARE structures and a starting structure taken from the initial ensemble. Fig. 3 *C* shows the RMSD (9.3 ± 1.2 A) of thermal fluctuations in the re-assembled ensemble. The *agreement* between Fig. 3 *B* and 3 *C* indicates that the RMSD between the starting and ending ensembles is essentially the same as that due to thermal fluctuations within the ending ensemble itself. The *difference* between Fig. *3 A* and *3 B* suggests that the re-assembled ensemble is not yet in equilibrium at the end of the re-assembly simulation; it is likely that there are slower processes that would bring the ensemble back to a state with lower thermal fluctuation. Thus, we may conclude that the CG model successfully re-assembles the SNARE structure into an ensemble close to but not completely the same as the starting one.

**FIGURE 3.**
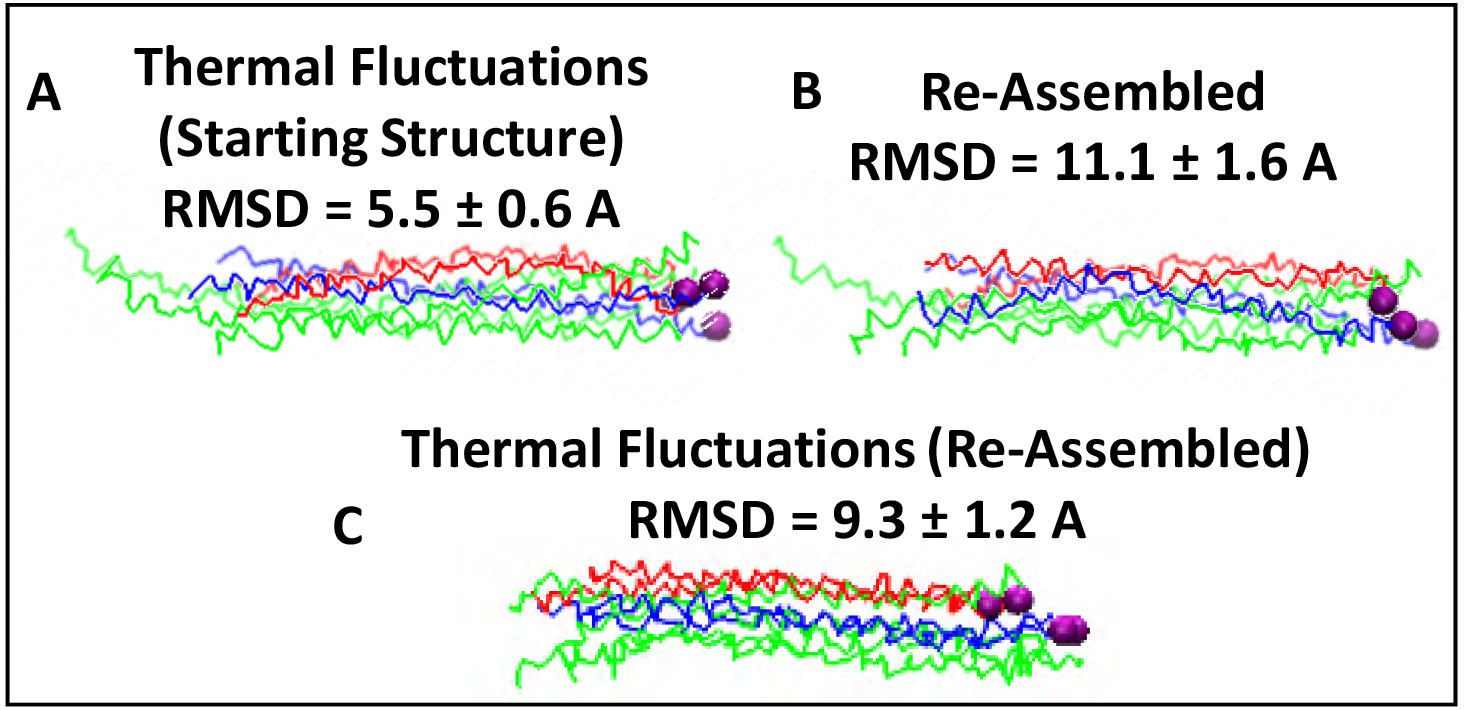
(*A*) RMSD of thermal fluctuations during a 1 µs trajectory of the starting SNARE structure without external force. (*B,C*) Reassembled SNARE bundle for the case corresponding toy Fig. 2 *A.* The structure is overlaid on the initial SNARE bundle structure for the disassembly simulation (*lighter colors*). (*B*) shows RMSD of the re-assembled structure with respect to one of the starting SNARE structures. (C) shows RMSD of the re-assembled structure with respect to one of the ending structures, i.e., due to thermal fluctuations in the re-assembled state. These results indicate that the ensemble of re-assembled structures is quite similar to the starting one. However, its fluctuations are significantly larger than those of the starting SNARE ensemble.

### 3.3 Assembly Time Depends Exponentially on the initial Separation

In order to determine the correlation between SNARE assembly time and initial Syb-Syx C-terminal separation, the assembly times from Fig. 2 were analyzed. The SNARE assembly time from 17 simulations is shown as a function of the initial Syb-Syx C-terminal distance in Fig. 4. Our results show that assembly time of partially assembled SNARE helices is quite rapid (hundreds of ns).

**FIGURE 4.**
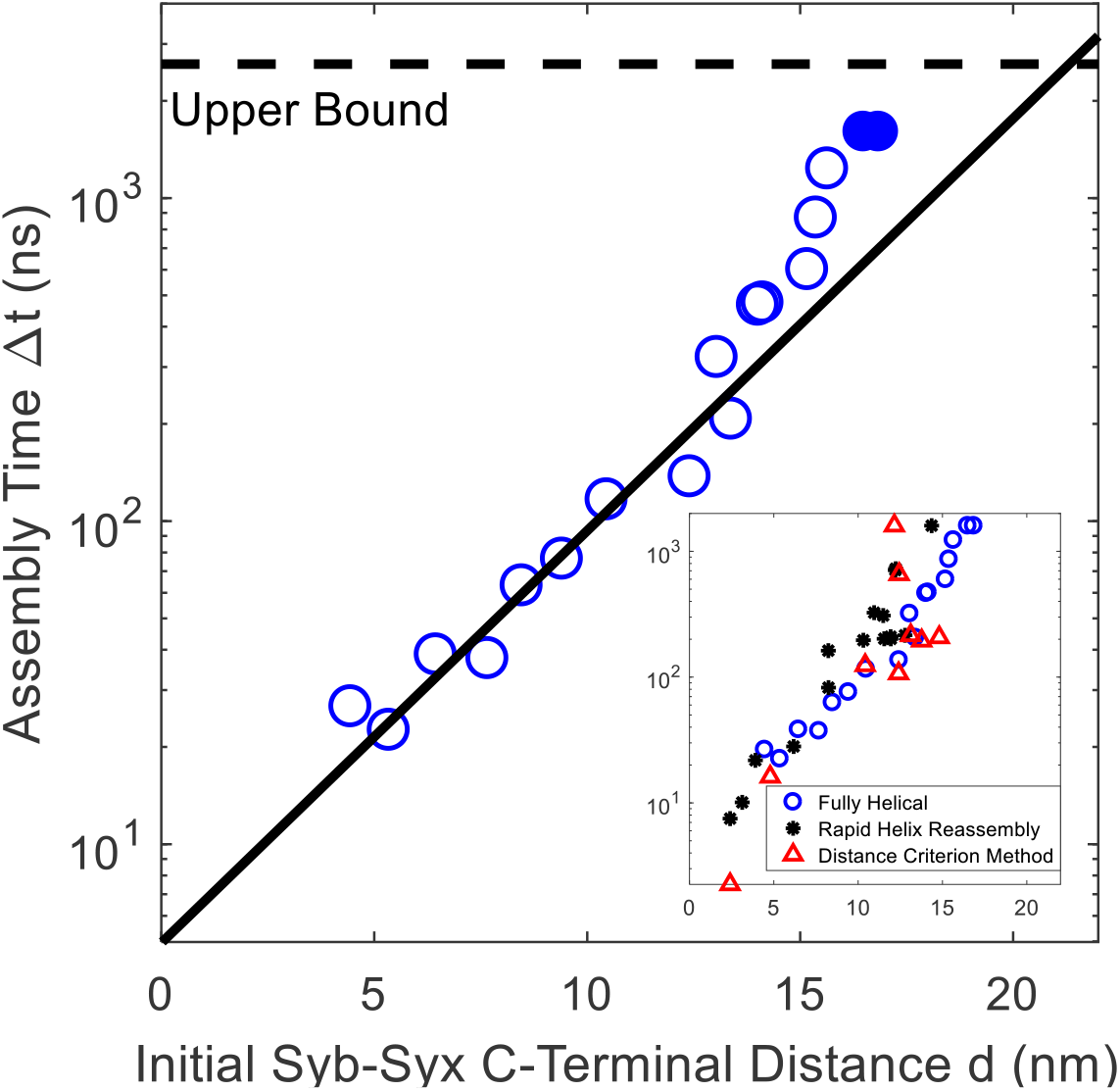
Growth of assembly time, **Δ*t***, as a function of initial Syb-Syx C-terminal distance, ***d***, was fitted with an exponential function, 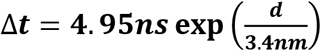. Of the 17 simulations shown, those with the lowest 10 values of initial separation all begin assembly immediately upon release of the pulling force. The remaining 7 cases (those with the 7 largest initial separations), upon release of the force, first fluctuate randomly at about a fixed separation and begin assembly once the N-terminals meet in partial assembly. In the simulations with the largest values of initial separation (filled circles), Syb did not assemble onto the t-SNARE within the duration of the simulation. (Inset) Assembly time (not accounting for assembly of helical turns) in simulations where Syb is allowed to melt during unzippering. Upon force release Syb re-forms as a helix following two different methods. Results show that overall assembly time is broadly unaffected, indicating robustness of our conclusions.

For separations less than about 15 A, there is strong growth in assembly time with increase in Syb-Syx C-terminal opening prior to start of the re-assembly simulation. For this range of initial separation, the results can be fit well by an exponential function of initial Syb-Syx C-terminal distance with a characteristic distance of 3.4 nm. For initial separation > 15A, the assembly time departs significantly from the exponential fit. As shown in Fig. S12 of Supporting Material, for the largest seven initial separations, there is a considerable period of time in the re-assembly simulation during which Syb fluctuates thermally before initiating contact with the t-SNARE bundle. This analysis of assembly time again indicates that rapidity of assembly is strongly enhanced by partial assembly.

### 3.4 Scaling model for helical Syb assembly onto t-SNARE

In order to better understand the time-duration for Syb re-assembly, we estimate characteristic relaxation times for two processes. In the first process, we estimate the characteristic time it takes for a randomly fluctuating cylinder to align with the substrate (Fig 5 *A*). In the second, we analyze the relaxation of a bent beam, representing Syb, onto a substrate representing the t-SNARE. The model reveals how assembly time depends upon physical properties and parameters and supports our interpretation of CG results.

**FIGURE 5.**
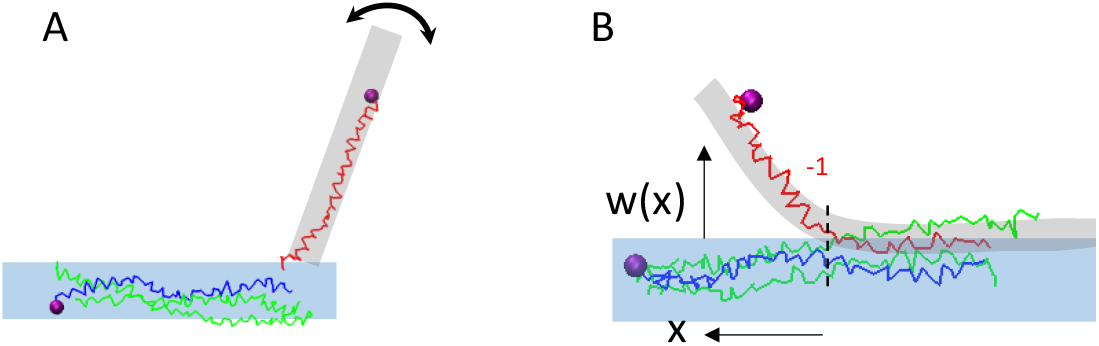
The assembly process for the syb helix can be thought of comprising two steps (*A*) rotational random walk of Syb until its N-terminus finds an orientation and location to initiate zippering, and (*B*) zippering driven by beam bending energy and adhesive interactions and resisted by viscous drag. The characteristic times associated with these two processes are consistent with those we find through BD simulations of the CG model.

#### 3.4.1 Estimation of Assembly Time for Small Initial Unzippering

We assume that inertia is negligible. The assembly process is driven by elastic energy in the bent Syb beam, and the rate of assembly is set by viscous dissipation. This is consistent with the fact that vibrational modes of this beam have time scales below 1 ns, determined by a normal mode analysis (NMA) or single SNARE helix fluctuations in Fortoul et al. (12), whereas assembly time is in the hundreds of ns to µs. This is also consistent with our assumption of solving the problem using Brownian Dynamics, not Langevin or molecular dynamics. Finally, the spectrum of fluctuations based on this assumption matches very well the MD results (12), indicating that the system is overdamped.

At any point, part of the beam has adhered to the substrate and part of it is out of contact. Let the length of the non-contacting region be *a*. The governing equation for the deflection of the beam (21) *w(x,t)*, is

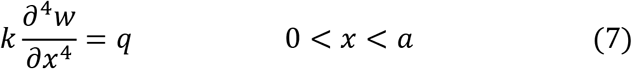

where *k* is the bending rigidity and *q* is the distributed load on the beam. This distributed load comes from the fluid’s viscous resistance as the beam travels through it. The beam, Syb, can be modeled as an ellipsoid where the radius of the major axis representing half the length of the non-contacting region, *a*/2, is about 4.5 nm and the radius of the minor axis representing the radius of the beam, *b*, is about 0.6 nm. The friction factor for such an ellipsoid moving sideways is given by (22)

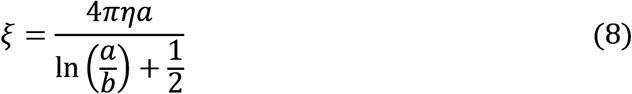

where *η* is the viscosity of water (8.9*x*10^−4^*Pa·s*). With these values of *a*, *b*, and *η* Eq. 8 gives 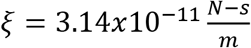. The distributed load *q* (*N/m*) is then given as the product of friction factor and local velocity, divided by the length of the ellipsoid:

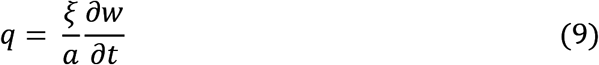

Using separation of variables (*w*(*x,t*) = *w*_*x*_(*x*)*w*_*t*_(*t*) (see Supporting Material for details) we get the solution of eq. (7) for fixed beam length as

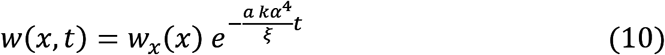

This immediately provides us with a characteristic relaxation time

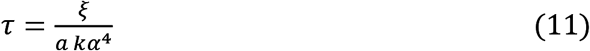

To determine the characteristic time, we need values for *k* and *α*. The bending rigidity, *k*, can be extracted from the frequency of the first normal mode of the beam, Syb, which is given as (23)

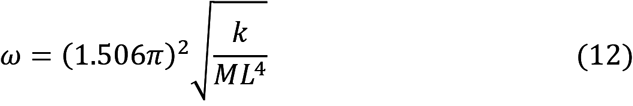

where *M* is the mass per unit length 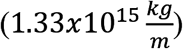 and *L* is the length of the beam (9 *nm*). Using the value for *ω* obtained from molecular simulation for Syb (2.22 *ns*^−1^) (12), we find that *k* = 8.58*x*10^−32^*Nm*^2^. The separation constant, *α*, is determined by imposing boundary conditions at the free end of the beam. It takes on one of a series of values corresponding to a set of mode shapes, *αa* = 1.875, 4.694, …. Any initial condition, i.e., the shape of the beam at *t=0*, can be represented as a sum of individual modes but all the higher modes decay quicker than the first mode so the characteristic decay time is based on the latter. The first mode of this solution can be combined with Eq. 12 to estimate the characteristic time

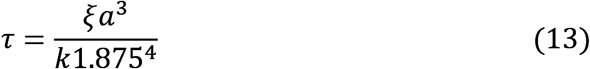

(For further details see the Supporting Material.) For SNARE structures with an initial unzippered state up to the ionic layer, *a*~4.5 *nm* (the length of helical Syb not in contact with the bundle). For such a separation, the time required for SNARE to assembly can be estimated, using Eq. 13, to be 2.5 *µs*. This timescale should be regarded as an upper bound for the assembly times found in Fig. 4, since we have neglected the influence of adhesive interactions, which will further speed up the process. This scaling analysis supports the conclusion from CG modeling that a helical Syb starting in contact with the rest of the SNARE can complete assembly well within physiologically relevant time duration.

#### 3.4.2 Estimation of Assembly Time for Syb Alignment

Next we estimate the time it would take for a randomly fluctuating rod, representing *Syb,* to align itself with the t-SNARE given an initially unfavorable orientation, such as structure E in Fig. 2. Again, the beam can be geometrically modelled as an ellipsoid. In order for assembly to occur, the ellipsoid must rotate itself to realign with the bundle. In order to determine the corresponding characteristic relaxation time, we use results for rotational diffusion of a prolate ellipsoid (24). Define *ρ* = 2*b*/*a*. The relaxation time is given by

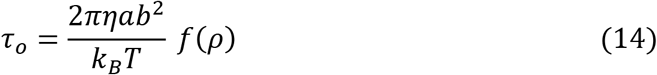

where *f*(*ρ*) was provided originally by Perrin (24, 25). Numerically, eq. (14) gives a timescale of ~100 *ns* which can be regarded approximately as the time it would usually take for a partially assembled Syb to find the correct orientation to initiate full assembly.

The sum of these estimated values can be interpreted as an upper bound for characteristic assembly time, which is consistent with the results shown in Fig. 4, supporting the interpretation that a partially assembled helical Syb will assemble with t-SNARE within a few microseconds and this period is governed by the time required for Syb to orient itself properly followed by assembly driven by beam elasticity and adhesive forces.

### 3.5 Effect of melting during unzippering

The results presented so far establish that the Syb helix assembles rapidly with t-SNARE to form the full SNARE bundle. In all the CG results presented so far, Syb remained in helical form during unzippering. In order to examine the sensitivity of our conclusions to this assumption, we also conducted assembly studies in which Syb was allowed to melt (following a force-based MC method) during its initial separation from the SNARE bundle, with subsequent re-formation of the helix in two different ways: (1) rapid Syb helix formation on force release, and (2) self-templated helix assembly. These are described in detail in the Supporting Material. In the rapid helix assembly model, the Syb helix quickly reforms when unzippering forces are released. In the self-templated helix assembly model, parts of Syb helix form helical turns during assembly as amino acids come close to the SNARE bundle (see Supporting Material for details). Results for these two cases, shown in Figs. S6 and S8 are qualitatively in good agreement with the results shown in Fig. 2. This is even though the initial state before assembly in these two cases was very different, in that the melting of Syb is considered in one and ignored in the other. Evidently, by re-formation of the Syb helix, the system loses memory of how it was unzippered. As before, there is a range of initial displacements that allows for quick and easy assembly, then a range that has a transition from slow to fast assembly, and finally for sufficiently large initial opening SNARE no longer assembles within the time frame of the simulation.

## 4 Discussion

We used a CG model and beam analysis to investigate the assembly of Syb onto t-SNARE. In particular, we addressed the large discrepancy between *in vitro* versus *in vivo* estimates of time required for SNARE assembly. Single molecule experiments performed *in vitro* demonstrated that SNARE assembly times are on the order of hundreds of milliseconds (10). Since the *in vivo* release process in nerve terminals occurs at a sub-millisecond time-scale it is unlikely that the final stages of exocytosis involve a full assembly of the SNARE bundle, starting from the fully unraveled state of Syb. Specifically, in this work we have shown that if Syb is in helical form and primed onto t-SNARE, it should assemble rapidly, at a microsecond time-scale. This is consistent with the idea that the discrepancy between *in vitro* and *in vivo* assembly times are due to the action of chaperones (in the latter case) that help to template the Syb into helical form and to bring it in sufficiently close proximity to t-SNARE.

That is, our study shows that zippering of a partially assembled SNARE complex is sufficiently fast to accommodate evoked synaptic transmission. In a recent study by Baker et al. (1, 26) it was proposed that Munc18, a chaperone, serves as a template for SNARE assembly. This crystallographic study showed that the yeast analog of Munc18 binds to a half zippered SNARE with the analog of Syb being in a partially helical form. This result suggests that Munc18 might serve as a template to assemble the prefusion SNARE complex with zippered N-terminus and helical Syb. Our study demonstrates that such a molecular mechanism could accelerate the final SNARE assembly from seconds to hundreds of nanoseconds.

Other details of our CG simulations are consistent with experimental findings. For example, numerous studies have reported on the significance of certain SNARE bundle layers. In an experimental study by Wiederhold et al. (8), layers −4 to −2 were determined to be the trigger site of the SNARE coiled coil bundle formation. When two residues within these layers were mutated (Syb^N49A, V50A^), SNARE assembly in solution was greatly compromised. In a second mutation (Syb^I45A, M46A^), this effect was even more prominent. Consistent with these studies, our simulations, demonstrate the importance of both layers −4 and −2 for the assembly process. During unzippering simulations, we find that Syb is strongly bound to t-SNARE at layer −4 particularly when melting models are used. The ionic layer (layer 0) has been determined experimentally to be critical to SNARE’s function (7, 9), since salt bridges are formed within this layer, which stabilize a half zippered confirmation of the SNARE bundle. In our simulations we observe that although SNARE assembly is a relatively slow process, it accelerates significantly once the SNARE complex is zippered to the vicinity of the ionic layer (layers −2 to 2), completing remaining assembly within ~100 ns.

Our CG model represents each residue by a single bead, thereby retaining chemical specificity. It includes intra-helical and inter-helical interactions and has been modified from a previous version to allow for force-based MC melting of Syb. CG simulations are complemented by scaling analysis of two components of the assembly process – random rotational diffusion of Syb to seek alignment with t-SNARE and subsequent un-bending of Syb as it assembles into the full SNARE bundle. We show that an upper bound estimate for the time required for these two steps is consistent with the CG results. As such, the model could serve as a computational tool to guide targeted mutagenesis and manipulate SNARE assembly.

## 5 AUTHOR CONTRIBUTIONS

NF & AJ developed the CG SNARE model; NF wrote the CG SNARE model code; NF, AJ and MB analyzed and interpreted results, and wrote the manuscript.

## 6 ACKNOWLEDGEMENTS

This study is supported by the NIH grant R01 MH099557. For MD computations, we acknowledge the support of the Extreme Science and Engineering Discovery Environment (XSEDE), which is supported by National Science Foundation grant number OCI-1053575, and specifically the Texas Advanced Computing Center (TACC) under grant number TG-MCB100049 and the National Institute of Computational Science (NICS). VMD/NAMD software is developed with NIH support by the Theoretical and Computational Biophysics group at the Beckman Institute, University of Illinois at Urbana-Champaign.

